# Single-Cell Transcriptomic Profiling Unveils Critical Metabolic Alterations and Signatures in Progression of Atherosclerosis

**DOI:** 10.1101/2024.02.06.579210

**Authors:** Limei Ma, Ying Zhao, Peng Xiang, Qingqiu Chen, Moustapha Hassan, Chao Yu

## Abstract

Metabolic dysregulation is recognized as a fundamental characteristic of cardiovascular diseases (CVDs), therefore mining metabolic patterns in these diseases would help to identify the possible pathogenic mechanisms and potential intervention targets. Atherosclerosis (AS), serving as the foundational pathology in numerous CVDs, represents a paramount global health concern. However, a systematic integrated analysis of the metabolic networks of AS is still lacking. In this study, we investigated and integrated single-cell RNA sequencing datasets from calcified atherosclerotic core (AC) plaques and patient-matched proximal adjacent (PA) portions of carotid artery tissue to generate metabolic flux profiling at single-cell level. Using scFEA and scWGCNA analyses, we discerned common metabolic changes in endothelial cells (ECs), myeloid cells, and smooth muscle cells. These altered metabolic modules were predominantly enriched in glucose-related pathways and were predicted to potentially facilitate metabolic bypass. Of particular interest, we observed an enrichment of metabolites produced from glucose/sialic acid metabolism pathway in both ECs and myeloid cells. This observation may partially account for their positive involvement in plaque formation, as previously discovered. Furthermore, we predicted that metabolic genes such as *HK1*, *ENO1*, *PFKL*, *LDHA*, *PGK1,* and *NANS* may be implicated in the detected metabolic flux disorder during AS progression. Additionally, we uncovered interactions between various cell types at single-cell level using CellChat. We noted heightened interactions between endothelial/SMC as well as myeloid/ECs in AC group, with ITGB2/VCAM and CD44/CD74 receptors potentially participating in these interactions, thereby fostering the development of pro-inflammatory plaque microenvironment. In conclusion, our study unveils metabolic shifts at single-cell level and identifies key gene signatures associated with metabolic disorders and cell-cell communication in atherosclerosis.

## Introduction

Atherosclerosis (AS) is a chronic disease characterized by low-grade, chronic inflammation of the arterial wall, and recognized as the leading cause of death and low quality of life worldwide [1]. Different compelling hypotheses have been proposed and studies have been performed to investigate the pathophysiology of atherosclerotic lesion formation and other cardiac complications such as myocardial infarction and stroke [2]. Despite these advances, the understanding of multiple key factors in AS procession, such as plaque heterogeneity, metabolic dysregulation and metabolic checkpoint, is still very limited [3].

Numerous studies have highlighted that metabolic dysregulation plays an important role in disease progression. Metabolic dysregulation encompasses deviations from normal metabolic processes in organisms , including disorders in carbohydrate, fat, and protein metabolism, and so on [4]. Atherosclerosis is characterized by lipid deposition, calcification, and angiogenesis within plaques [5]. Recent investigations propose that the metabolic reprogramming of glucose, cholesterol, fatty acid, and amino acid in cells such as macrophages, endothelial cells (ECs), and vascular smooth muscle cells (VSMCs) significantly contributes to inflammation at all stages of atherosclerosis, from lesion initiation to advanced stages, and even to lesion regression [6–9]. For example, disruptions in glycolysis, along with the accumulation of its metabolite lactic acid, was reported to regulate endothelial inflammation and the formation of plaques [10]. Additionally, the infiltration of immune cells, such as macrophages, can also induce alterations in immunometabolism within plaques [11]. Consequently, these studies emphasize the critical contribution of metabolic disturbances in the progression of AS. However, to achieve a systematic view of metabolic flux alterations in the AS process remains challenging due to the high heterogeneity and complicated cell-cell interaction within AS plaques.

Single-cell RNA sequencing (scRNA-seq) technology enables research at unprecedented resolution by analyzing global gene expression and regulation pathway alternation in different cell populations during disease progression. Nowadays, scRNA-seq has emerged as a powerful tool for investigating cell trajectory, RNA velocity and cell-cell communication [12, 13]. ScRNA-seq has already been employed to determine cell heterogeneity in AS pathological conditions. In addition, scRNA-seq together with a graphneural network-based method were used to evaluate metabolic fluxes (scFEA), to illustrate the cellular metabolic diversity and predicate metabolic gene signatures in diseases development [14]. Guo et al have utilized scRNA-seq/scFEA analysis to comprehensively assess metabolic shifts at single-cell level, and found that the metabolites like pyruvate and fumarate decreased in ECs during stroke and Alzheimer’s disease progression [4]. To better understand metabolic dysregulation in AS development, there is an urgent need for a consensus single-cell reference to discern metabolic shifts in various cell types.

In this study, we investigated and integrated two single-cell RNA sequencing (scRNA-seq) data profiles, comprising calcified atherosclerotic lesions (referred to as AC) and patient-matched proximal adjacent artery tissue (referred to as PA). Our analytical approach included scFEA analysis, scWGCNA, cell-cell communication analysis, and Gene Ontology (GO) analysis to delineate metabolic flux alterations and identify relevant cell types with their corresponding effector genes. Our findings revealed an up-regulation of glycolysis, hexosamine, and sialic acid metabolites across various cell types, particularly in myeloid cells and ECs. Additionally, specific genes such as *HK1, ENO1, PFKL, LDHA, PGK1, and NANS*, were recognized as potential markers involved in the regulation of glycolysis to sialic acid metabolism disorder. These genes were also implicated in promoting the formation of a pro-inflammatory microenvironment. Furthermore, our findings also indicated the enhanced interactions between endothelial/SMCand myeloid/ECs in AC group. Notably, receptors such as ITGB2/VCAM and CD44/CD74 were identified as potential regulators in these intracellular interactions, contributing to the formation of an inflammatory microenvironment. In summary, this comprehensive map of metabolic flux in human atherosclerosis represents a pivotal advancement in translating mechanistic knowledge, laying the foundation for the development of new targeted therapies.

## Materials and methods

### Study design and data collection/integration

A flow diagram summarizing the entire study design is provided in Fig. 1A. To conduct a comprehensive analysis of atherosclerotic core (AC) plaques and patient-matched proximal adjacent (PA) portions of carotid artery tissue, we acquired high-dimensional datasets from the Gene Expression Omnibus (GEO, https://www.ncbi.nlm.nih.gov/geo/). This included single-cell RNA sequencing (scRNA-seq) data from patients undergoing carotid endarterectomy, comprising 3 AC and 3 PA samples, via accession number GSE159677. Additionally, we integrated dataset GSE155512, enhancing the diversity and depth of our dataset [15, 16]. Single-cell sequencing dataset preprocessing and analysis The above dataset was analyzed using Seurat package (v4.1.0), including filtering, normalization, dimensionality reduction, clustering, and identification of cell types. Specifically, cells expressing fewer than 600 genes or more than 4,000 genes were excluded to eliminate potential low-quality “doublets.” Additionally, a cap of 20,000 UMIs was set, aligning with established standards for filtering low-quality cells [17]. Cells with over 10% mitochondrial genes (MT-) were also removed to ensure data purity. To identify and eliminate potential doublet cells, we also employed DoubletFinder (v2.0.3; [18]). DecontX within the celda package (v1.6.1; [19]) addressed ambient RNA contamination, a common issue in single-cell RNA-seq. Following critical filtering, the data underwent normalization for downstream analysis.

**Figure 1.**
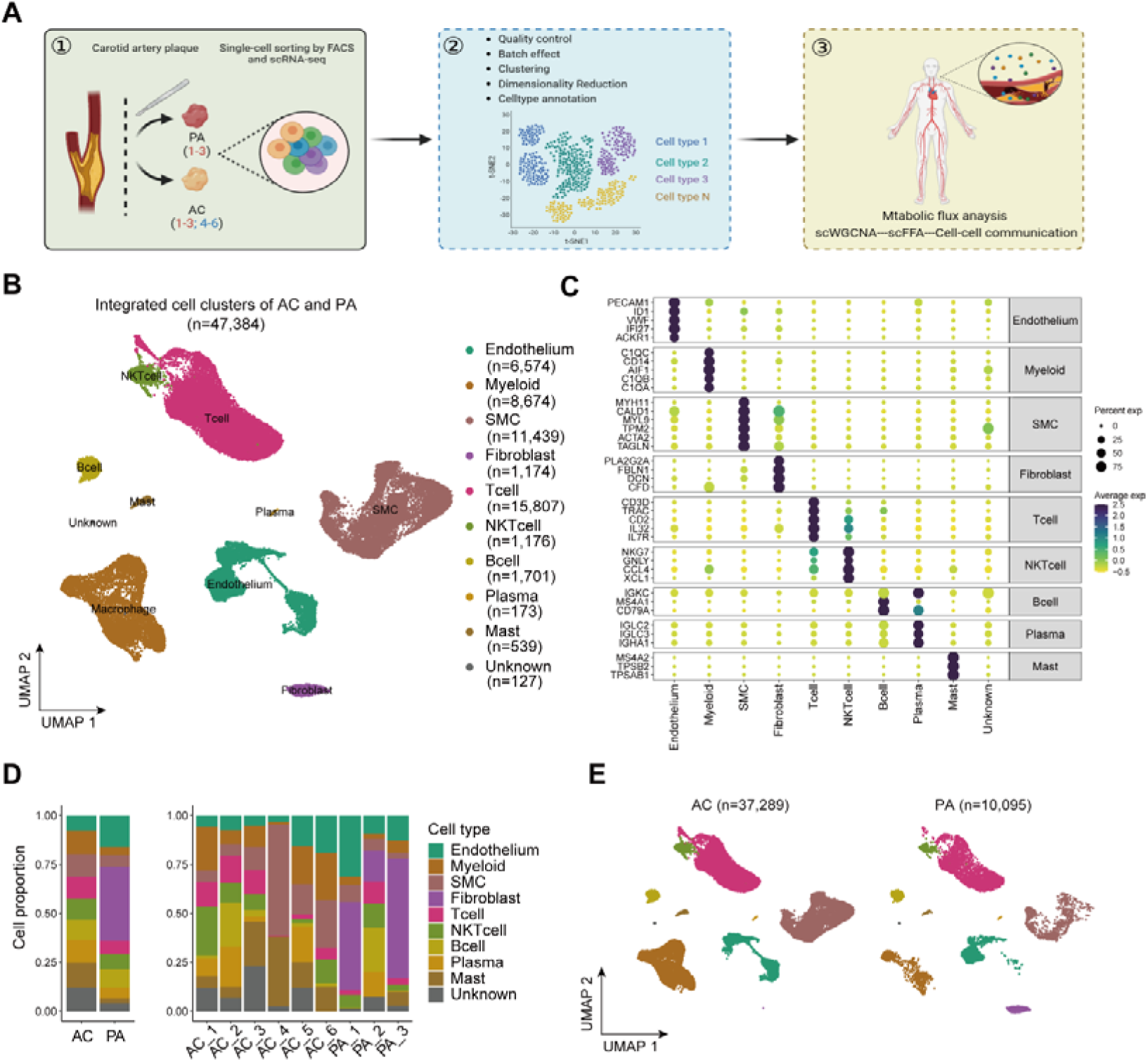
Single-cell atlas of human atherosclerotic coronary arteries. (A) Flow diagram of the the study design. PA, Patient-matched proximal adjacent portions of carotid artery tissue aortic dissection. (B) UMAP plots of all cells, with cells colored by (left to right): 10 major cell types. (C) Dot plot showing the top markers of each cell type. Dot size corresponds to the proportion of cells within the group expressing each gene, and dot color corresponds to its expression level. (D) The number of cells of each cell type, and the corresponding patients. (E) UMAP regional occupancy analysis demonstrating relative changes in cell density comparing atherosclerotic plaque to healthy controls. Atherosclerosis-associated cell states are revealed by increased local abundance of cells in regions of the UMAP plot (log_2_FC, log2 fold change). AC, calcified atherosclerotic core plaques; FACS, fluorescence-activated cell sorting; scRNA-seq, Single-cell RNA-sequencing; scWGCNA, Single-cell weighted gene co-expression network analysis; AS, atherosclerosis; HUVECs, human umbilical vein ECs; GSE159677, AC plaques and PA portions of carotid artery were collected from 3 patients and named as PA1-3, AC1-3; GSE155512, carotid arteries were obtained from 3 patients, and named as AC4-6.

Standard procedures for data normalization and scaling, as well as for comprehensive cell analysis, were shown in responsible Figs as well as in Fig. S1, S2. We also pursued data integration using Seurat’s FindIntegrationAnchors function and the Harmony package (v1.2.0) for batch effect correction, ensuring the harmonization of data across different samples. In the dimensionality reduction process, classic PCA and the t-SNE algorithm were employed. From the log-transformed expression data, 2,000 highly variable genes were selected for principal component analysis (PCA). Subsequently, 30 principal components (PC1 to PC30) were chosen for shared nearest neighbor (SNN) algorithm and t-distributed stochastic neighbor embedding (t-SNE) analysis, with a resolution of 0.8 for cell clustering.

For the identification of different cell types, we sourced the differential expressed genes (also called marker genes) of each cluster through “FindAllMarkers” function. Further, these genes were used for cellular identification. Also, we applied the marker genes which are specifically expressed in various cells to determine the cell type in each cluster. Then the clusters were merged by cell type, and the differential expressed genes of each cell type were rediscovered for subsequent analysis.

### Construction of a single-cell metabolic flux profile

The acquisition of single-cell metabolic flux profiles primarily involved the application of a graph neural network algorithm known as single-cell flux estimation analysis (scFEA, v1.1.2) [14]. The 168 metabolic modules used in this study were directly imported from the algorithm’s official Github page (https://github.com/changwn/scFEA). [4]. Initially, we re-clustered and dimensionally reduced the single-cell expression matrices of metabolic modules using the Seurat package (v4.1.0), followed by visualizing the single-cell distribution of flux values with FeaturePlot function. Cell populations were then segmented based on cell and disease types, with the exclusion of groups containing fewer than 200 cells and generated metabolic expression profiles. To identify flux values that differed between AC and PA under specific cell type conditions, we screened for changes through differential values. The Wilcoxon rank-sum test was employed to statistically assess these changes, identifying significant metabolic alterations (*P* value < 0.05, after adjustment).

### Single-cell weighted gene co-expression network analysis (scWGCNA)

For network-based analysis, we employed high-definition Weighted Gene Co-expression Network Analysis (hdWGCNA, v0.2.24). This method facilitated the exploration of co-expression patterns among genes, unveiling potential regulatory networks and key driver genes associated with AS. In the integration of single-cell data into metacells, we set specific criteria, including a minimum threshold of 50 cells per metacell, a maximum sharing limit of 10 cells between metacells, and a nearest-neighbors parameter of 25, which ensured the specificity of our analysis. Focusing on the top 2000 variable genes, we constructed a signed adjacency matrix to identify gene modules, setting the SoftPower parameter at 18 to balance sensitivity and specificity. Average hierarchical clustering was employed for gene tree and eigengene clustering. We utilized the ModuleCorrelogram function to visually represent the correlation of co-expressed modules. The FindDMEs function was utilized to identify differential co-expression modules between AC and PA. Additionally, the ModuleNetworkPlot function was employed to generate gene network diagrams specifically for modules of interest.

Establishment of genes signatures related to glucose/hexosamine/sialic acid metabolism Genes associated with glycolysis and TCA cycle were sorted based on the previous report [20]. Briefly, genes encoding key enzymes of glucose metabolism were selected including 14 genes (HK1, HK2, HK3, PFKL, ALDOA, GAPDH, PGK1, PGAM1, ENO1, ENO2, ENO3, PKM, LDHA, PFKFB3). And 7 genes (GFPT1, GFPT2, GNPNAT1, PGM3, UAP1, OGT, OGA) were selected into the hexosamine pathway as previous study reported [21]. Also, genes including GNE, NANS, NANP, CMAS, CMAHP, NPL, ST3GAL1-6, ST6GAL1-2, ST6GALNAC1-6, ST8SIA1-6,

NEU1-4 were selected as maker genes of sialic acid metabolism pathway according to the previous reports [22]. Then the above genes were integrated and regarded as the glucose/hexosamine/sialic acid metabolism gene signature, and then used to identify subgroups of metabolism activity. The detailed information about the above genes were shown in Table S1.

### Cell-cell communication analysis

Cell-cell communication analysis was performed using CellChat (v1.6.1) on a set of variable genes derived from all cells. And we considered interactions such as myeloid cells and ECs. contacts. Interaction networks and ligand-receptor pairs were identified for all cell groups and selected cell subtypes in both AC and PA states, with default parameters applied. Subsequent differential analysis was carried out. For visualization, the netVisual_diffInteraction function was utilized to depict differential network diagrams between AC and PA states. Additionally, the netVisual_circle function was employed to create comparative network diagrams of cell interactions, retaining the top 30% of interactions. To assess pathway signal strength differences between AC and PA states, the rankNet function was applied. Moreover, the netVisual_bubble function was used to illustrate the action intensity of ligand-receptor pairs in both AC and PA states.

### Ontology Enrichment Analysis

Gene Ontology (GO) enrichment analysis provided a comprehensive understanding of the biological functions, processes, and pathways enriched in AS progression. Therefore, we performed GO enrichment analysis using the hypergeometric test and the clusterProfiler package (v3.18.1). Specifically, we performed enrichment analysis for endothelial cell subtypes, Myeloid subtypes’ marker genes, and co-expressed module genes to discern their biological significance. The outcomes of the analysis were effectively visualized using the dotplot function.

### Statistical Analysis and Visualization

All statistical and visualization analyses were carried out using R software (v4.0.5). The Wilcoxon rank-sum test and Kruskal-Wallis test were applied as appropriate. *p*-values > 0.05 were considered not statistically significant and denoted as n.s., while *p*-values less than or equal to 0.05 were represented as follows: \**p* ≤ 0.05, \*\**p* ≤ 0.01, \*\*\**p* ≤ 0.001, and \*\*\*\**p* ≤ 0.0001. *p*-values were adjusted based on Bonferroni correction. Visualization of the results was accomplished using ggplot2 (v3.3.6), ggpubr (v0.4.0), ggrepel (v0.9.1), and ComplexHeatmap (v2.6.2) for the generation of high-quality graphics.

## Results

### Single-cell data integration and revealed cellular heterogeneity

Development of atherosclerosis (AS) and the process of plaque formation involve the participation of various cell types [23]. In this study, we initiated an investigation into the distinctions among different cell types during AS development by integrating two single-cell sequencing datasets (GSE155512 and GSE159677). The workflow for data processing was illustrated in Fig. 1A. Following quality control and filtering (Fig. S1A), we identified 47,384 cells from calcified atherosclerotic core plaques (AC group) and patient-matched proximal adjacent aortic portions (PA group) for subsequent single-cell analysis. Subsequently, we applied uniform manifold approximation and projection (UMAP) and categorized these cells into nine cell types (Fig. 1B) by comparing gene expressing profiling with previously reported specific markers for each cell type, as depicted in Fig. 1C.

Additionally, we observed a reduction in the proportions of ECs and SMC in the AC group compared to the PA group (Fig. 1D). This observation suggests that dysfunction in these cells significantly contributes to the formation of calcification plaques and inflammation of the aortic wall, as previously confirmed [24]. Furthermore, the AC group exhibited a notable increase in the proportion of macrophages, T cells, NK cells, and mast cells, indicating a substantial infiltration of adaptive immune cells during pathogenesis associated with plaque formation. To better illustrate the differences between cell types during AS progression, we also focused on regions of the UMAP plot where the local cell density increased more than 2-fold in AS disease compared to control group. As depicted in Fig. 1E, we observed significant alterations in the proportions of cell types such as ECs, macrophages, and SMCs, underscoring the contribution of cellular heterogeneity to the AS process.

### scWGCNA identified perturbed co-expression modules linked to metabolic dysfunction

Subsequently, we employed scWGCNA, a novel systems biological method known for delineating essential modules and pathways in diseases, to elucidate the intricate relationship between genes expression alteration and phenotype difference during the progression of AS [25]. We utilized highly variable genes (HGVs) identified through the FindMarkers function in the Seurat Package, and succeeded to discover nine distinct co-expression modules, as illustrated in Fig. 2A, Fig. S2A, Fig. S2B. When focusing on the average expression level of genes in each cell, we observed that modules 9 and 6 predominantly comprised in ECs, while modules 4, 5, 7, and 8 were predominantly associated with myeloid cells. Modules 2 and 3 exhibited enrichments of genes associated with SMCs, and module 1 featured genes indicative of T/B cells (Fig. 2B). Additionally, modules 2/3 displayed a significant positive correlation with modules 4/6/8, and modules 6/9 exhibited a positive correlation with modules 5/8, while modules 4/5/7/8 showed a positive correlation with modules 6/9, as illustrated in Fig. 2C. This suggests a strong involvement of myeloid cells in communication with ECs and SMCs during the AS process. Moreover, we observed that modules 4/5/7/8 exhibited a significant increase in the AC group, while modules 6/9 displayed a decrease compared to the PA group (Fig. 2D). This indicates that the infiltration of inflammatory cells such as macrophages and the disruption of endothelial homeostasis positively contribute to the exacerbation of the AS process.

**Figure 2.**
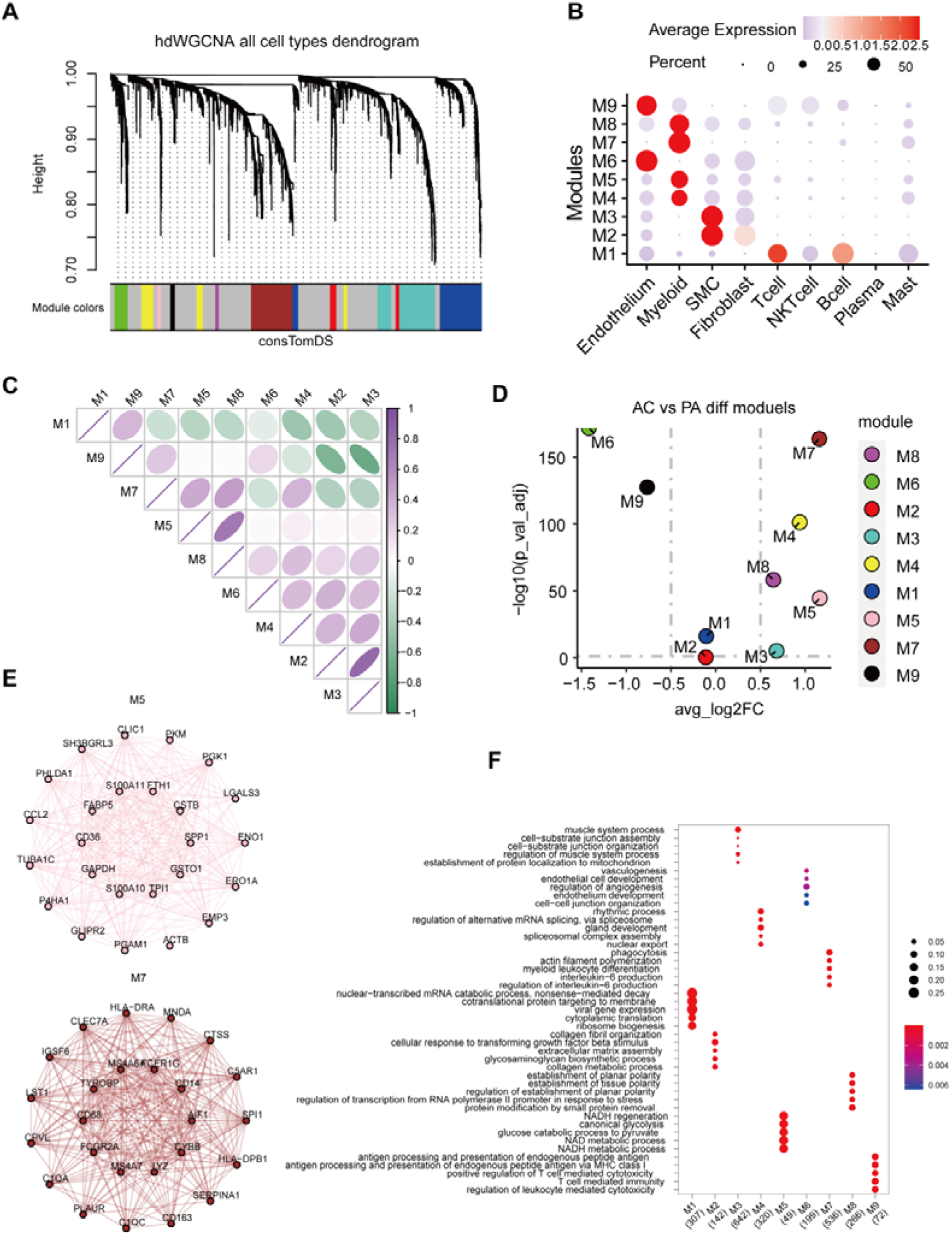
scWGCNA revealed that the differences of cluster and possible pathway. (A) Highly variable genes were clustered into 9 modules through hdWGCNA. (B) Average expression of genes in different modules and its association with cell types. (C) The correlations between different modules. (D) Dot plot of the different module scores in AC vs PA. (E) Gene-gene interaction analysis in module 5 and 7. (F) Go analysis revealed the enrichment of pathways in different modules.

Furthermore, we conducted gene-gene interaction analysis, revealing notable correlation between cells, particularly in modules 5. There was a significant enrichment of metabolic genes such as ALDOA, ENO1, GAPDH, GPI, LDHA, PGAM1, and PGK1, which are highly implicated in the glucose metabolism pathway. In addition, modules 7 featured several genes including GLUL, PFKFB3, NANS, ST3GAL1, and ST8SIA4, which were reported to be associated with the sialic acid metabolism pathway (Fig. 2E, Fig. S2C). Moreover, GO analysis highlighted a significant enrichment of genes associated with critical metabolic processes, including NADH metabolic process, glucose metabolism, and the glycolysis pathway in modules 5. And genes associated with the secretion of inflammatory factors, such as IL-6, were enriched in modules 7 (Fig. 2F). This intriguing observation suggested that myeloid cells may play a significant role during plaque formation where an alternation in glucose metabolism occurs. And the sialic acid pathway appears to be linked to the pro-inflammatory state of myeloid cells in AS process.

### scFEA analysis revealed distinct metabolic change across various cell types

To comprehensively understand pivotal metabolic signatures during the progression of AS, we employed an integrated algorithm named single-cell flux estimation analysis (scFEA) [14]. This approach allowed us to systematically evaluate 168 metabolic modules and 70 metabolites at the single-cell level during AS progression. Following quality control and filtering (Fig. S1B), we observed an enhanced heterogeneity of metabolic modules across several cell types, including ECs, myeloid cells, and SMCs during AS progression (Fig. 3A, B). The analysis of metabolic genes from 168 metabolic modules confirmed that modules indicative of myeloid cells, SMCs, and ECs exhibited the highest difference in altered genes (Fig. S2D). Due to changes in cell proportion, the fluxes of metabolic modules in fibroblasts from PA group were not able to be caulculated. Therefore, we focused on ECs, myeloid cells, and SMCs in the follow-up analysis.

**Figure 3.**
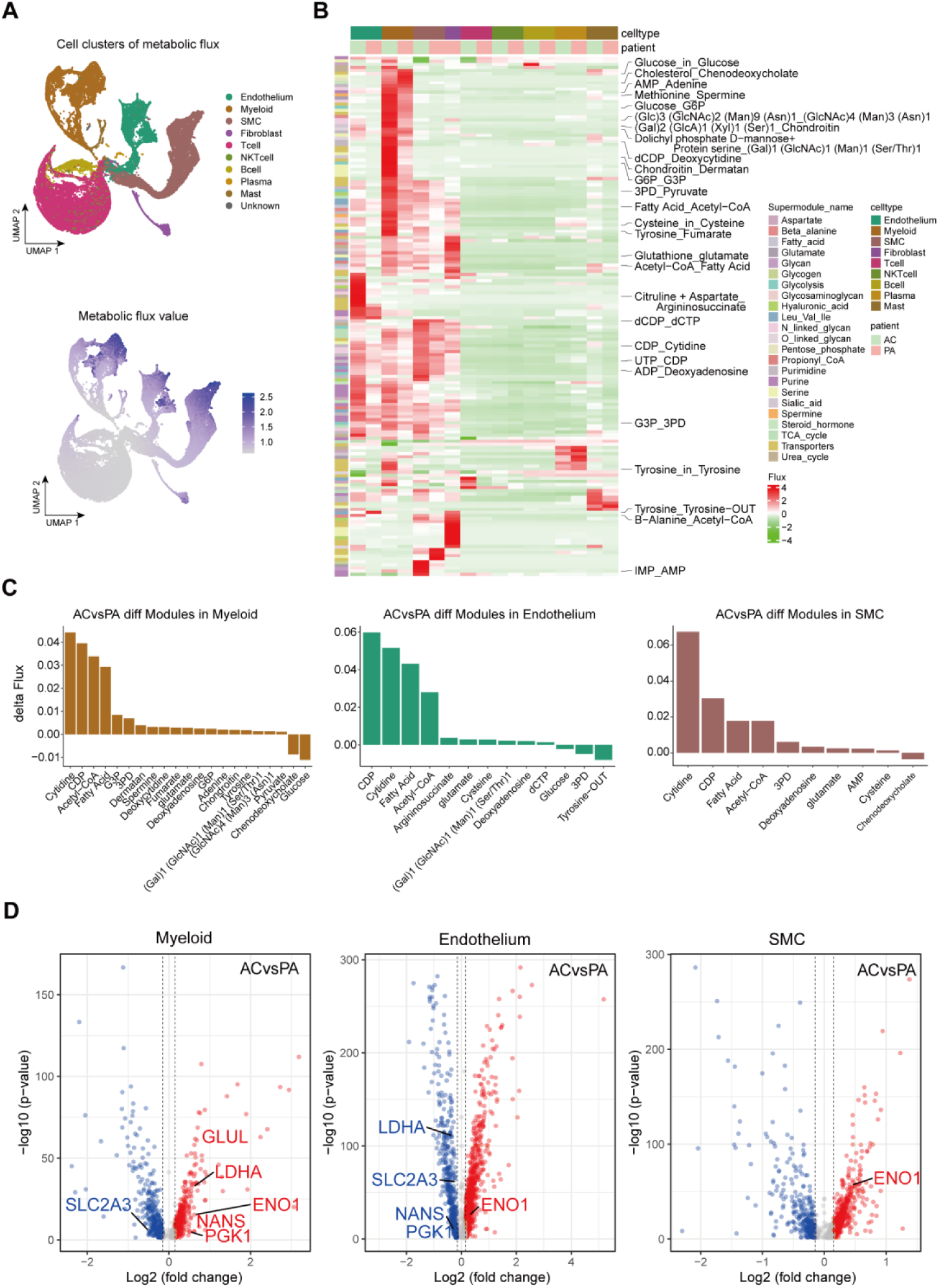
Single-cell metabolic flux mapping reveals the heterogeneity of metabolism between cell types. (A) The metabolic flux in different cell clusters using UMAP plots. (B) Profiles of predicted fluxome of 168 metabolic modules., each row on the y-axis displays the flux of metabolic module for all cells under various group (shown on the x-axis). (C) Top 10-20 metabolites predicted to increase or decrease in AC groups compared to PA group. (D) Volcano map revealing differences in enzyme expression in ECs, myeloid cells and SMCin AS group when compared with AC group. The vertical axis provides average log2FC values, and the horizontal axis presents the difference between proportions. The color of each dot represents the direction of change. Enzymes that differ significantly are highlighted on the figure.

In ECs, we observed a simultaneous increase in the fluxes of several metabolic modules, including M-4 (3PD to Pyruvate), M-15 (3PD to Serine), M-34 (Acetyl-CoA to Fatty Acid), M-107 (Glucose-6-phosphate to UDP-N-acetylglucosamine), M-108 (Glucose-6-phosphate to Glucose-1-phosphate), M-110 (UDP-glucuronic acid+UDP-N-acetylglucosamine to Hyaluronic acid) and M-123 ((GlcNAc)4 (Man)3 (Asn)1_(Gal)2 (GlcNAc)4 (LFuc)1 (Man)3 (Neu5Ac)2 (Asn)1). These alternations indicated that glucose metabolism in ECs might dominate over other pathways including lipid metabolism and post-translational modification. In addition, the fluxes of metabolic modules in myeloid cells also displayed an obviously enhancement, such as M-2 (G6P to G3P), M-16 (Serine to Pyruvate), M-107 (Glucose-6-phosphate to UDP-N-acetylglucosamine), M-112 (UDP-N-acetylglucosamine to CMP-N-acetylneuraminate), M-123 (GlcNAc)4 (Man)3 (Asn)1 to (GlcNAc)7 (Man)3 (Asn)1), M-2 (G6P to G3P), M-131 (Chondroitin to Dermatan), which suggested that the enhancement of glucose associated side metabolism in myeloid cells. Furthermore, we also noticed similar alternations in modules including M-4, M-15, m-34, M-107, M-108 from SMC cells (Fig. 3B, Table. S2). Collectively, these results suggest that ECs, myeloid cells, and SMC went through metabolism dysregulation under AS pathological conditions.

Next, we assessed the alterations in metabolite levels to further explore the metabolic disorders. Fig. 3C indicated that ECs, myeloid cells and SMC had the similar feature in the alternation of several key metabolites . Especially, the levels of cytidine, cytidine 5’-diphosphate (CDP), fatty acid, acetyl-CoA were increased, and the level of glucose was decreased at the same time. Cytidine as well as CDP, triphosphate of cytidine, are endogenous metabolites in pyrimidine pathway, and has been reported in regulating lipid disorder [26, 27]. In addition, acetyl-CoA is known to be produced through glucose and acetate uptake pathway, serving as the essential building block for fatty acid and isoprenoid biosynthesis [28, 29]. From the present results, we concluded that the metabolic flux may switch from glucose metabolism to fatty acid oxidation and lipids formation. Furthermore, we also found that the intermediate metabolites from hexosamine/sialic acid pathway displayed a visible up-regulation, including (GlcNAc)4 (Man)3 (Asn)1, (Gal)1 (GlcNAc)1 (Man)1 (Ser/Thr)1, and chondroitin, dermatan. Especially, these products were strongly increased in myeloid cells. Taken together, we identified that the enhancement of glucose towards its ancillary metabolic pathways, such as the sialic acid pathway in myeloid cells, might play a crucial role in AS progression as confirmed before [30, 31].

Additionally, we conducted an analysis of the enzymes from 168 metabolic modules, to identify their expression level changes in AS models. As shown in Fig. 3D, enzymes such as GLUL, LDHA, ENO1, PGK1, NANS were significantly increased in myeloid cells, which partly supported the results from scWGCNA analysis and also indicated these enzymes might be responsible for the alternations of related metabolites. Interestingly, we also found that the level of ENO1 was increased in both ECs and SMC, indicating that ENO1 might be highly involved in the metabolic shift during AS progression. Also, we observed that the expression of solute carrier family 2 member 3 (SLC2A3), encoding the predominantly neuronal glucose transporter 3 (GLUT3), was decreased in myeloid cells and ECs from AC group. Collectively, these findings provide additional evidence that AS plaques serve as a central hub for metabolic dysregulation. Notably, the dysregulation of myeloid cells, such as macrophages, appears to play a pivotal role in the progression of AS.

### scFEA analysis highlighted glycolysis/hexosamine/sialic acid metabolic shift in plaque

Reprogramming of glucose metabolism was considered as main emerging hallmark of cardiovascular disease [32]. Specially, our present results have indicated that the intermediate metabolites from glycolysis/hexosamine/sialic acid metabolism displayed an obvious increase in myeloid cells and ECs, indicating that dysfunction of glucose to sialic acid metabolism might play essential roles in maintaining the homeostasis of those cells. Moreover, sialic acid is a component of cell surface sugar moieties, and accumulation of sialic acid in plasma has been reported to be highly associated with AS progression [33].

Several studies have investigated sialic acid levels in individuals with AS or those at high risk of AS [30, 31, 33–39]. The results showed that sialic acid levels in patients with heart failure, acute coronary syndrome, coronary atherosclerosis, type 2 diabetes mellitus and hyperlipidemic were higher than the relative control group (Table 1), further confirmed that sialic acid could be a potential metabolic maker to indicate the disease progression as we reported previously. Moreover, our previous study indicated that N-acetylneuraminic acid (Neu5Ac), the most widespread form of sialic acid, was increased in plaque and was positively correlated with AS progression [30]. Therefore, we hypothesis that the activation of sialic acid metabolism pathway in myeloid cells and ECs might partly be responsible for the accumulation of sialic acid in plaque, and further contribute to the formation of plaque microenvironment, eventually promote AS progression.

**Table 1.**
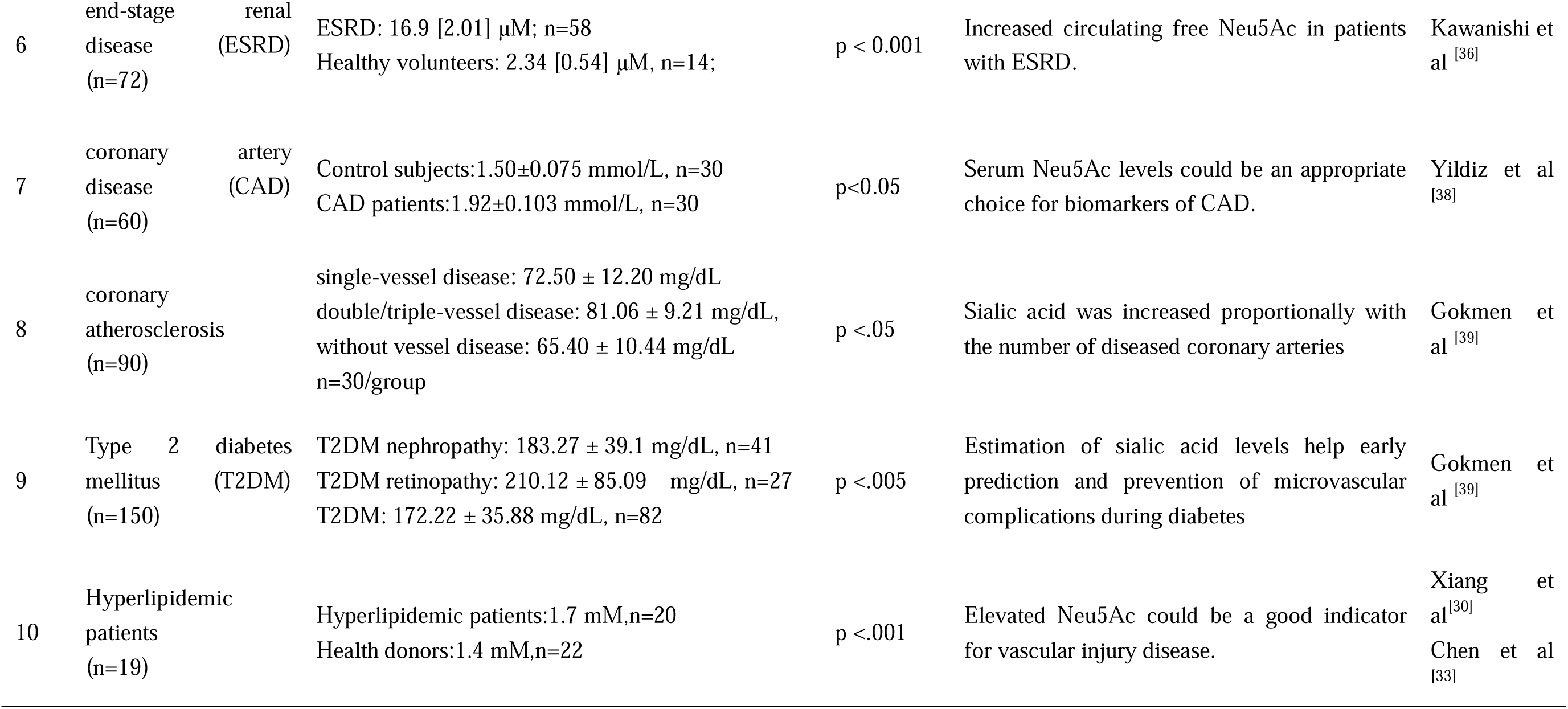
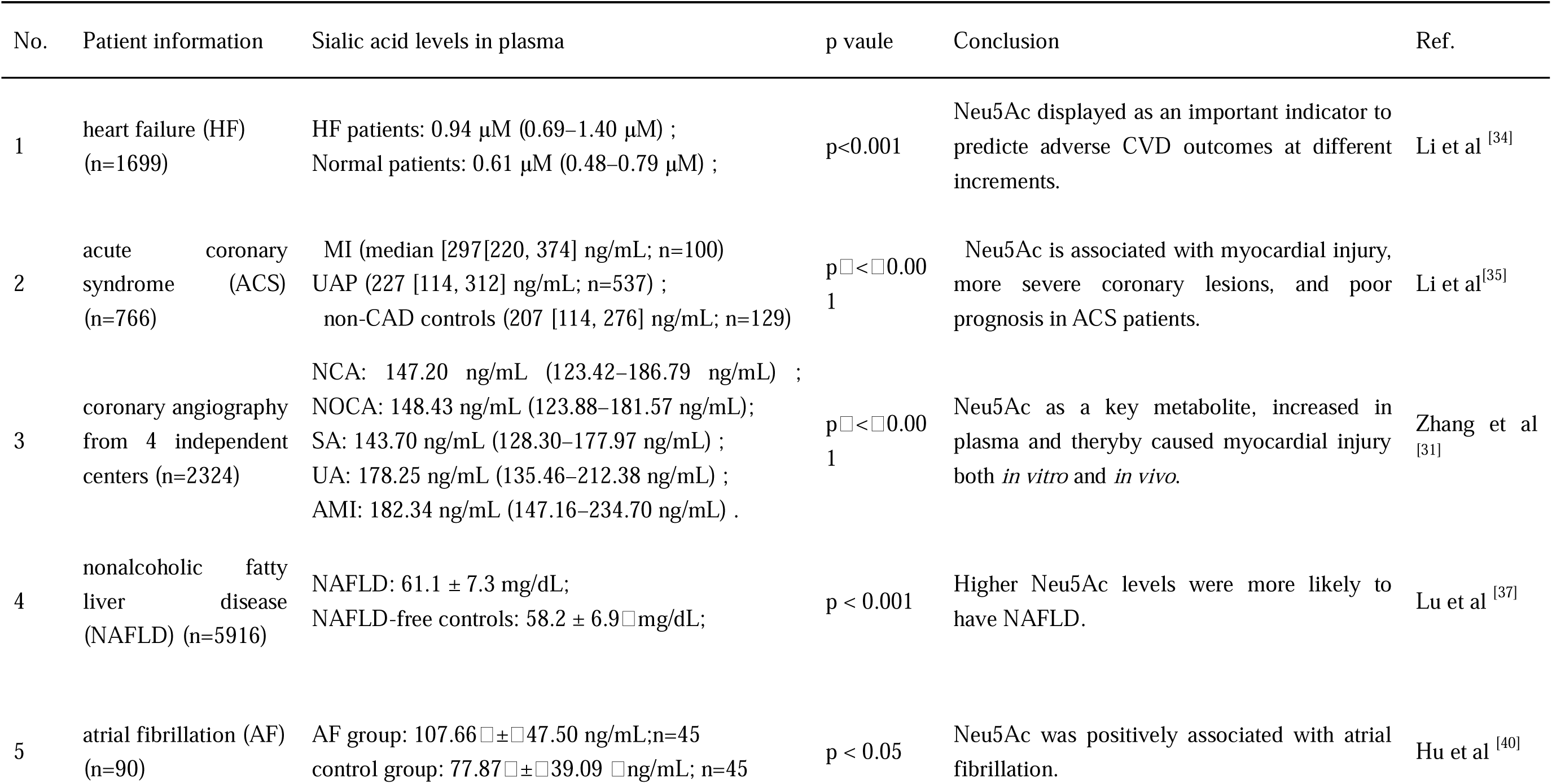
Prognostic Value of Elevated Serum sialic acid in patients with cardiovascular diseases (CVD)

To further investigate the underlying mechanism upon sialic acid metabolism activation, we compared the detailed metabolites from glycolysis/hexosamine/sialic acid pathway in different cell types. As shown in Fig. 4A, myeloid cells, ECs and SMC displayed as the most metabolically active cell types. It is worth noting that the metabolites in myeloid cells were mostly enriched, and metabolic shifts including glucose to glucose 6-phosphat, then metabolized to UDP-N-acetylmannosamine, and further metabolized into CMP-N-acetylneuraminate were strongly enhanced in myeloid cells and ECs, and particularly higher in myeloid cells from AC group (Fig. 4B). Furthermore, we investigated the expression of genes associated with glycolysis, hexosamine and sialic acid pathway between PA and AC group. The genes of interest were selected based on the molecular signatures database and previously published results[20]. As shown in Fig. 4C, D, HK1, ENO1, PGK1, PFKL, and LDHA exhibited significant up-regulation in myeloid cells from AC group. And only ENO1 displayed increasing level in ECs. In addition, genes involved in sialic acid devo metabolism for example NANS, NEU1 were significantly higher in myeloid cells, and the expression of ST6Gal1 was moderately increased in ECs. These results suggested that these genes might be responsible for the metabolic dysfunction in regulating the metabolic shift in AS plaque. Since sialic acid anabolism occurs often in myeloid cells , increasing of NANS might be involved in the regulation of sialic acid accumulation in plaque. On the other side, the ECs were mainly dominated by sialic acid transport metabolism, which was also confirmed from the results in Fig. 3B that metabolites for sialylation modification were specially increased in ECs.

**Figure 4.**
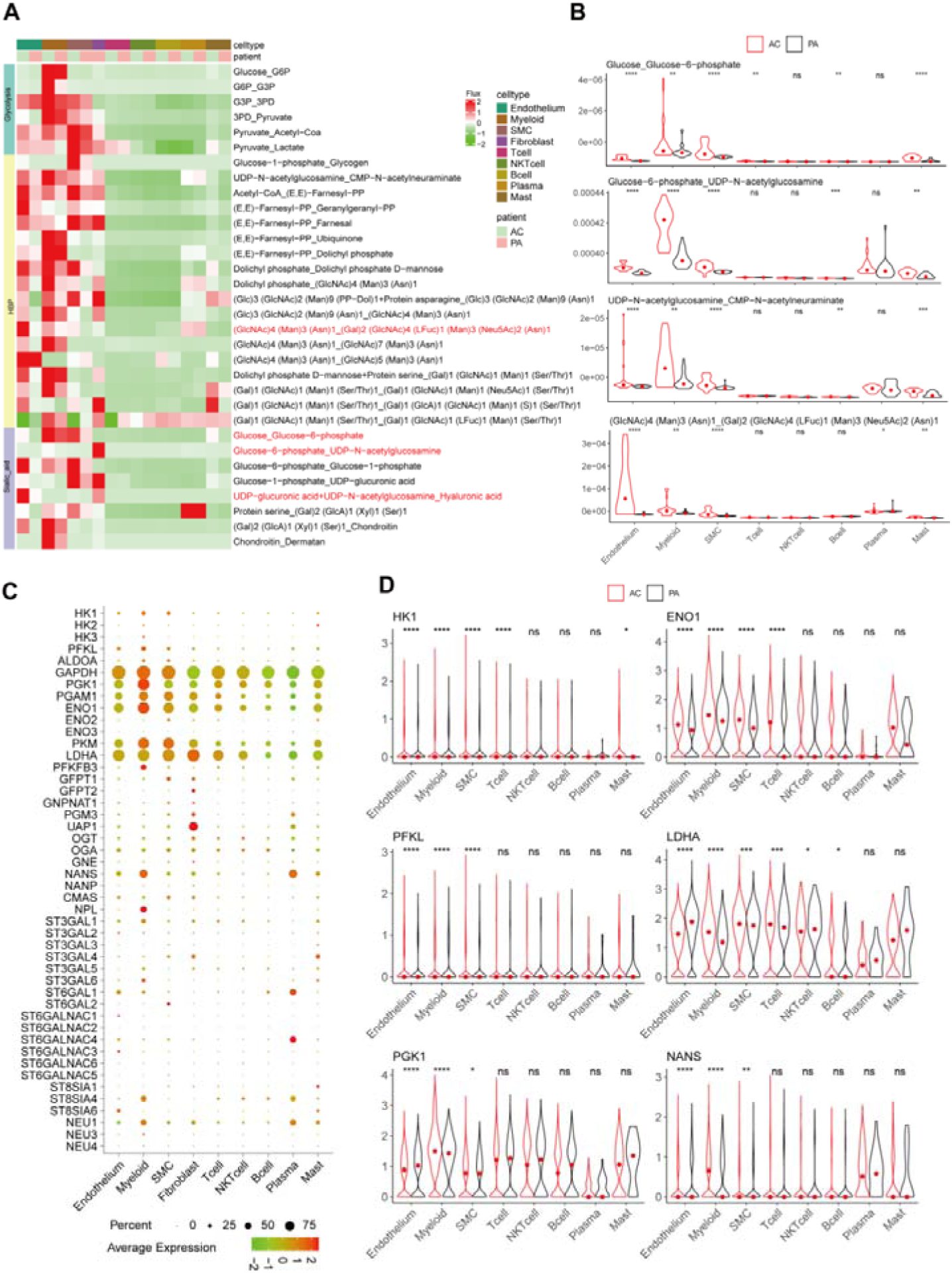
Single-cell metabolic flux mapping showed metabolic shift from glucose to sialic acid metabolism in different cell types and diseases. (A) Profiles of the predicted fluxome of glycolysis/hexosamine/sialic acid metabolic modules. Each row of y-axis displays the flux of metabolic module from cells under various conditions (shown on the x-axis). (B) Violin plots showing classical metabolic intermediates of glycolysis/hexosamine/sialic acid pathway in different cell types. The vertical axis provides average log_2_FC values, and the horizontal axis presents the difference between metabolites in AC group when compared to the control group. (C) Dot plot showing the genes signatures associated with glycolysis/hexosamine/sialic acid metabolic pathway. Dot size corresponds to proportion of cells within the group expressing each gene, and dot color corresponds to expression level. (D) The average expression of genes like HK1, ENO1, PFKL, LDHA, PGK1 and NANS in different cell types. \**p* < 0.05; \*\**p*< 0.01; \*\*\**p*< 0.001.

### Cell–cell communications revealed correlations between myeloid cells and ECs

It has been suggested that metabolic dysfunction can lead to the release of pro-inflammatory cytokines, exacerbating the disruption of intercellular communication in atherosclerotic plaques [40]. Here, we further characterized discrepancies in the molecular interactions among distinct cell populations from spatial sites of lesions using CellChat. The network was refined using molecular markers to identify potential specific intracellular communication between cell populations. Eight major cell populations (SMCs, myeloid cells, ECs, T cells, NK cells, B cells, plasma cells, mast cells) exhibited complex interactions, with a higher number of interactions observed in the AC group (Fig. 5A, 5B). Notably, we observed distinct interactions between SMCs, ECs, and myeloid cells in lesions, with a significant increase in interactions in the AC group (Fig. 5C, D, Fig. S3A). Furthermore, differential signaling pathway enrichment analysis showed that osteopontin (SPP1)-mediated signaling pathways, TGF-β pathways and IL-1β pathways were strongly enriched in lesion samples (Figure 5E). In addition, we also performed a detailed analysis of changes in signaling-receptor levels for all important pathways. Some pathways were displayed actively in special cell types from lesion samples, such as SPP1 pathway, TGF-β and IL-16 receptor pathway were enriched in myeloid cells. And CD46, CD34, NOTCH, EPHB pathway were specially enriched in ECs (Fig. S3B). These results were consistent with previous reports implicating these pathways particularly contributed to the intercellular communication between endothelial and macrophages, and upregulated in AS progression as predicted [40, 41]. Moreover, we also noticed that HSPG pathway was enriched in AC group and receptors like CD44 as well as CD74 were shown to be increased in AC group (Fig, 5E). And tight junction makers like CDH5 was also enhanced in AC group, ICAM interaction with ITGB2 was displayed to be promoted within ECs and myeloid cells (Fig, 5F). These results further suggested that interaction between ECs and myeloid cells like macrophages should be considered as a contributor to plaque development. Additionally, several receptors including CD74, CD44, ICAM might participated in the process of ECs and macrophages interaction. However, the underlying mechanism still needs further investigation. Also, the link between the sialic acid dysfunction and cell-cell interaction, as well as the role of above receptors remain to be further explored.

**Figure 5.**
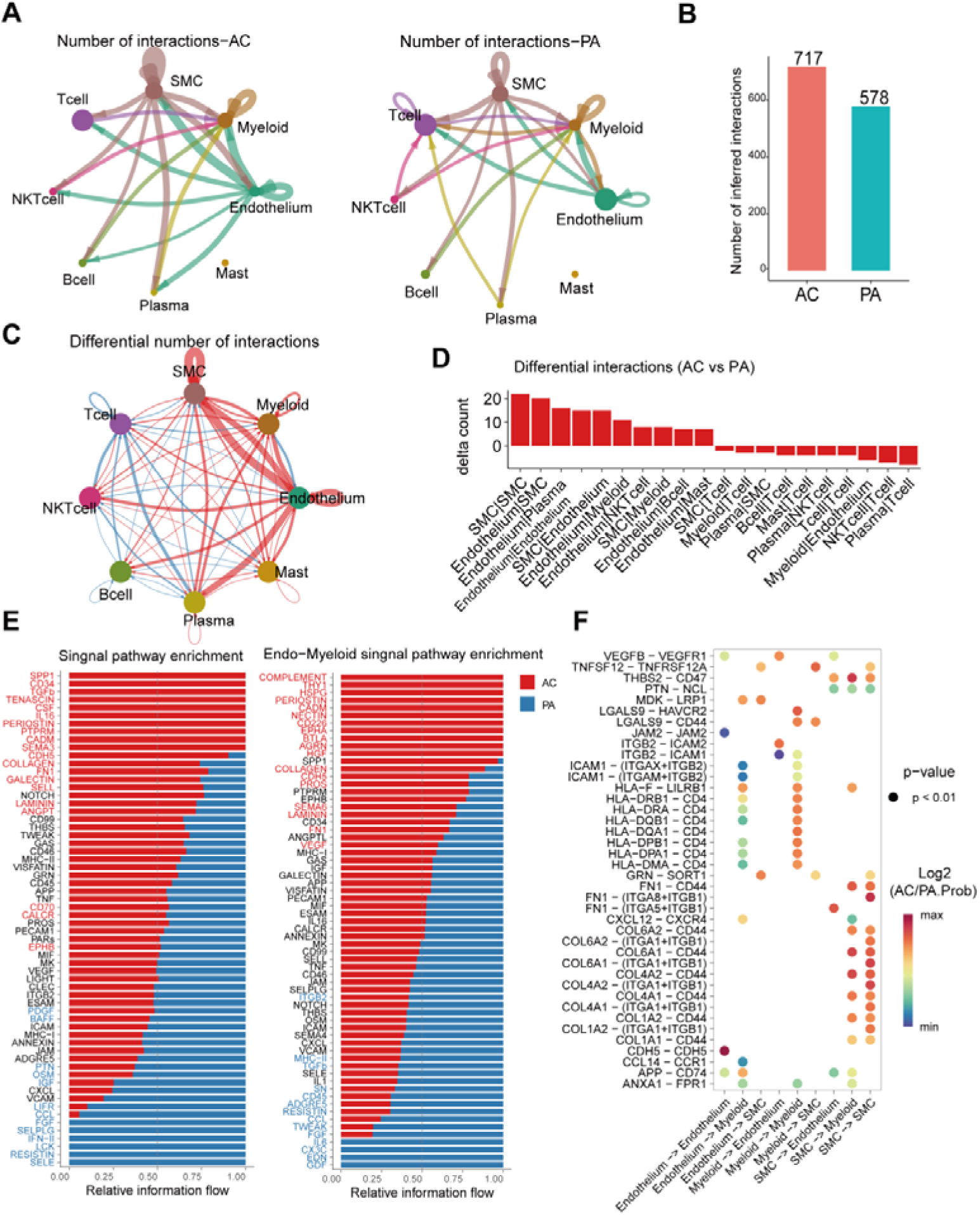
Intercellular communication in different cell types during atherosclerosis progression. (A, B) Overall intercellular communication between each cell population in PA/AC group and numbers of interaction. The color and width of bands correspond to numbers of ligand-receptor pairs. (C, D) Circle plot as well as top 10 different interaction showing the number changes between major cell types. Blue lines: decreased communication in AC; Red lines: increasing communication in AC. Line thickness: number of unique ligand-receptor interactions. (E) Significant signaling pathways ranked based on differences in the overall information between PA and AC groups. Red, top pathways enriched in AC; Blue, enriched in PA. (F) Dotplot showing the activity of the top receptor interaction between main cell types including ECs, myeloid cells, and SMC.

## Discussion

Metabolic dysfunction intricately links with CVDs, and atherosclerosis (AS) manifests as a consequence of a multistep metabolic process that ultimately leads to cardiovascular pathology [42]. Despite evidence showcasing cellular heterogeneity within AS plaques, the potential connections between altered metabolic pathways and cellular heterogeneity have not yet to be systematically investigated [43]. In this study, we conducted scRNA-seq of AS plaques and adjacent arterial tissue in patients. Leveraging scWGCNA, scFEA, and cell-cell interaction analyses, we confirmed the augmentation of glucose metabolism in AS group, particularly within distinct cell types including myeloid cells, ECs, and SMCs. Remarkably, we also observed an elevation in the branched metabolic pathway of glucose metabolism, specifically sialic acid pathway, characterized by increased levels of intermediate products and featured genes within myeloid cells. Furthermore, the identification of key genes, such as GLUL, LDHA, ENO1, SLC2A3, PGK1, and NANS, underscores their pivotal roles in regulating metabolic shifts during AS progression. Additionally, our study revealed that the interaction between myeloid cells and ECs positively contributes to the establishment of a pro-inflammatory plaque microenvironment. In summary, our present investigation not only advanced the understanding of the intricate association between cellular heterogeneity and metabolic dysfunction in AS process, but also underscored the critical role of sialic acid dysfunction in myeloid cells or ECs as a significant contributor to AS development.

In atherosclerosis research, single-cell studies have significantly advanced our understanding of cellular heterogeneity in vessels [44]. As described before, GSE159677 datasets was aimed to analyze the transcriptional profiles of six main cell populations and also identified key gene drivers of pathogenic biological processes in SMCs and ECs [16]. And original results from GSE155512 suggested that the trajectories of SMC trans differentiation during AS as well as identification of molecular targets [15]. Even though these two datasets focus on different cell types, they shared a similar patient feature. Therefore, we investigated and integrated these two datasets to further investigate the potential metabolic shift during AS progression. Consistent with the results from two datasets, we first confirmed the cellular heterogeneity of AS plaque. In addition, our results of proportions in plaque also suggested the decrease of ECs along with a significant increase of immune cells proportions in AC group. These results further confirmed the evidence that infiltration of adaptive immune cells contributes to plaque formation [41]. Moreover, our results replenished the local cell density changes in AS plaque, and confirmed the density alterations of ECs, macrophages, SMCs mainly contributing to AS heterogeneity. Together, our present study improved the previous study, and pointed out that metabolic dysfunction of cells including ECs, myeloid cells strongly involved in AS progression.

Glycolysis serves as a critical energy acquisition pathway for diverse cell types, including ECs, SMCs, macrophages, and others. Under pathological conditions, it plays a significant role in AS by influencing processes such as lipid deposition, calcification, and angiogenesis within plaques. Additionally, glycolysis produces lactic acid as a metabolite, contributing to the complex metabolic landscape associated with AS [5]. To our knowledge, there is no systematic research to explore the metabolic alternations at single cell level [3]. Here, we utilized scFEA to discern potential alterations in 168 metabolic modules and 70 metabolites from AS lesions. Our findings revealed an increase in metabolites, including cytidine, CDP, fatty acids, acetyl-CoA, and a decrease in glucose levels, observed across ECs, myeloid cells, and SMCs. As numerous literature reported, cytidine and CDP are endogenous metabolites in the pyrimidine metabolism pathway, and cytidine supplementation in ob/ob mice could alleviate certain aspects of dyslipidemia, improve hepatic steatosis as well [26]. Nevertheless, its potential impact on AS progression, as well as its interrelation with glycolysis, remains inadequately explored. Hence, our study partially elucidates this phenomenon, and its effect still needs to be further investigated. Moreover, acetyl-CoA was reported to be essential for oxidative metabolism of glucose and fatty acids. And high levels of acetyl-CoA in hepatocytes could stimulate lipid biosynthesis, ketone body production and the diversion of pyruvate metabolism towards gluconeogenesis [45]. Moreover, the balance between glucose consumption and fatty acids preservation could prevent the shift of metabolic flux into the anabolic pathway and maintains catabolic metabolism for energy production [46]. In the present study, we further confirmed the imbalance between acetyl-CoA, glucose and fatty acids at single cell level. These findings indicated that glucose metabolism might shunt to other metabolic branches, such as hexosamine/sialic acid pathway.

Sialic acid belongs to the neuraminic acid derivative family and originates from glucose to hexosamine pathway [30]. Epidemiological studies have revealed that the elevated plasma sialic acid especially Neu5Ac was highly associated with the increased risk of cardiovascular events [47]. And our previous results also pointed out the pro-atherosclerotic effect of sialic acid through inducing endothelial inflammatory injury or promoting macrophage M1 polarization [30, 33, 48]. However, the mechanism underlying the accumulation of sialic acid in AS plaque still needs further explore. In the current study, we investigated almost all the genes in the sialic acid regulatory pathways, as well as the related metabolites. From the results, we found that some metabolic modules including glucose-6-phosphate to UDP-N-acetylglucosamine, and then metabolized into CMP-N-acetylneuraminate, as well as chondroitin to dermatan were increased along with metabolites. This finding partly indicated that sialic acid metabolic flux dysfunction should originated from glycolysis pathway.

In addition, scFFA as well as scWGCNA analysis featured several genes including GLUL, LDHA, ENO1, PGK1 and NANS were mostly up regulated in AS lesions, especially in myeloid cells, which indicating that myeloid cells might be sialic acid dysregulation center in AS process. Macrophages are regarded as the main type of myeloid cells and inflammatory cells in atherosclerotic plaque [49]. New insights have shed light on the importance of metabolic and functional reprogramming in macrophages for AS progression [9]. Specially, researchers have linked the macrophage GLUL, LDHA, ENO1 as well as PGK1 to polarization, pro-inflammatory factors release and plaque vulnerability [50–53]. Our present study also pointed out that these genes are also involved in the regulation of metabolic shifts, and highly associated with inflammation. However, few studies explored the relation of NANS with vascular homeostasis as well as inflammatory regulation. Concerning its highly relevant with sialic acid metabolism, targeting NANS might also be a potential strategy to regulate sialic acid disorder during AS progression. Collectively, our present study confirmed the previous study, and featured the importance of above genes in regulating metabolic flux shift during AS progression.

Furthermore, our study revealed a complex network of interactions among different cell types, with increased interactions observed in VSMCs, myeloid cells, and ECs. Previous reports have already pointed out that the crosstalk between macrophages and ECs was predicted to increase in atherosclerosis. Interaction between ICAM/VCAM and ITGB2 (immune cells, especially inflammatory macrophages) was speculated to be essential for AS pathogenesis [40]. Here, we further confirmed the above conclusion, and found the junction protein, CDH5, was enhanced in AC group, which indicating that the crosstalk might induce injury of ECs as reported before [54]. Moreover, it have been found that high levels of CD74 could promote inflammatory responses and eventually contributed to ECs-macrophages interaction [55]. Additionally, CD44 build a link between Neu5Ac and atherosclerosis as Chen at al reported [56]. In conjunction with our findings highlighting the involvement of CD74 and CD44, these results further suggest that these surface proteins might establish a link between the crosstalk of macrophages/endothelial cells and the disorder in sialic acid metabolism during AS progression. However, a more in-depth exploration of the underlying mechanisms is warranted.

Taken together, our present study sheds light on dynamic cellular and metabolic changes in AS progression using scRNA-seq together with comprehensive systemic analysis including scWGCNA, scFFA and cell communication analysis. Here, we confirmed the cellular heterogeneity in AS plaque. In addition, we also identified metabolic flux shifts, especially glycolysis/hexosamine/sialic acid metabolism alternation in myeloid cells. Further, we featured several genes including GLUL, LDHA, ENO1, PGK1, NANS, were highly associated with those metabolic shifts. Moreover, cell-cell communication provided valuable insights into the intricate mechanisms underlying AS progression.

## Abbreviations

CVDs: Cardiovascular diseases
AS: Atherosclerosis
scRNA-seq: Single-cell RNA sequencing
scFEA: single-cell flux estimation analysis
ECs: ECs
VSMCs: vascular smooth muscle cells
GO: Gene Ontology
scWGCNA: single-cell weighted gene co-expression network analysis
PCA: principal component analysis
SNN: shared nearest neighbor
t-SNE: t-distributed stochastic neighbour embedding
GEO: gene expression omnibus
UMAP: uniform manifold approximation and projection
HGVs: highly variable genes
Neu5Ac: N-acetylneuraminic acid
AC: Atherosclerotic core
PA: Adjacent portion

## Acknowledgements

The authors would like to thank Dr. JunWei Song and Qinkuan Qin (Sichuan University) for their generously sharing bioinformatics experience and codes. Additionally, the authors would like to thank the data provider from GEO dataset.

## Author contributions

Limei Ma, Chao Yu conceived the study; Limei Ma performed most of the bioinformatics analysis and wrote the manuscript; Ying Zhao gave much suggestions on study design and help to revised the manuscript; Qingqiu Chen and Peng Xiang revised the manuscript and verified the data; Chao Yu and Moustapha Hassan supervised the research and helped manuscript editing.

## Availability of data and materials

The data used to support the findings of this study are available from GEO database under accession codes.

## Declarations

All authors have approved the contents of this paper. And We declare no competing interest relevant to this article.

## References

1. Bäck M, Yurdagul A, Tabas I, Öörni K, Kovanen PT. Inflammation and its resolution in atherosclerosis: mediators and therapeutic opportunities. Nat Rev Cardiol. 2019; 16: 389–406.

2. Tomaniak M, Katagiri Y, Modolo R, de Silva R, Khamis RY, Bourantas CV, et al. Vulnerable plaques and patients: state-of-the-art. Eur Heart J. 2020; 41: 2997–3004.

3. Libby P, Ridker PM, Hansson GK. Progress and challenges in translating the biology of atherosclerosis. Nature. 2011; 473: 317–25.

4. Guo G, Fan L, Yan Y, Xu Y, Deng Z, Tian M, et al. Shared metabolic shifts in ECs in stroke and Alzheimer’s disease revealed by integrated analysis. Sci Data. 2023; 10: 666.

5. Xu R, Yuan W, Wang Z. Advances in Glycolysis Metabolism of Atherosclerosis. J Cardiovasc Transl Res. 2023; 16: 476–90.

6. Tamargo IA, Baek KI, Kim Y, Park C, Jo H. Flow-induced reprogramming of ECs in atherosclerosis. Nat Rev Cardiol. 2023; 20: 738–53.

7. Shi J, Yang Y, Cheng A, Xu G, He F. Metabolism of vascular SMCin vascular diseases. Am J Physiol Heart Circ Physiol. 2020; 319: H613–H31.

8. Miano JM, Fisher EA, Majesky MW. Fate and State of Vascular SMCin Atherosclerosis. Circulation. 2021; 143: 2110–6.

9. Groh L, Keating ST, Joosten LAB, Netea MG, Riksen NP. Monocyte and macrophage immunometabolism in atherosclerosis. Semin Immunopathol. 2018; 40: 203–14.

10. van Tuijl J, Joosten LAB, Netea MG, Bekkering S, Riksen NP. Immunometabolism orchestrates training of innate immunity in atherosclerosis. Cardiovasc Res. 2019; 115: 1416–24.

11. Tabas I, Bornfeldt KE. Intracellular and Intercellular Aspects of Macrophage Immunometabolism in Atherosclerosis. Circ Res. 2020; 126: 1209–27.

12. Slenders L, Tessels DE, van der Laan SW, Pasterkamp G, Mokry M. The Applications of Single-Cell RNA Sequencing in Atherosclerotic Disease. Front Cardiovasc Med. 2022; 9: 826103.

13. Örd T, Lönnberg T, Nurminen V, Ravindran A, Niskanen H, Kiema M, et al. Dissecting the polygenic basis of atherosclerosis via disease-associated cell state signatures. Am J Hum Genet. 2023; 110: 722–40.

14. Alghamdi N, Chang W, Dang P, Lu X, Wan C, Gampala S, et al. A graph neural network model to estimate cell-wise metabolic flux using single-cell RNA-seq data. Genome Res. 2021; 31: 1867–84.

15. Pan H, Xue C, Auerbach BJ, Fan J, Bashore AC, Cui J, et al. Single-Cell Genomics Reveals a Novel Cell State During Smooth Muscle Cell Phenotypic Switching and Potential Therapeutic Targets for Atherosclerosis in Mouse and Human. Circulation. 2020; 142: 2060–75.

16. Alsaigh T, Evans D, Frankel D, Torkamani A. Decoding the transcriptome of calcified atherosclerotic plaque at single-cell resolution. Commun Biol. 2022; 5: 1084.

17. Luecken MD, Theis FJ. Current best practices in single-cell RNA-seq analysis: a tutorial. Mol Syst Biol. 2019; 15: e8746.

18. McGinnis CS, Murrow LM, Gartner ZJ. DoubletFinder: Doublet Detection in Single-Cell RNA Sequencing Data Using Artificial Nearest Neighbors. Cell Syst. 2019; 8.

19. Yang S, Corbett SE, Koga Y, Wang Z, Johnson WE, Yajima M, et al. Decontamination of ambient RNA in single-cell RNA-seq with DecontX. Genome Biol. 2020; 21: 57.

20. Shi J, Wu Z, Wu X, Huangfu L, Guo T, Cheng X, et al. Characterization of glycometabolism and tumor immune microenvironment for predicting clinical outcomes in gastric cancer. iScience. 2023; 26: 106214.

21. Willems AP, van Engelen BGM, Lefeber DJ. Genetic defects in the hexosamine and sialic acid biosynthesis pathway. Biochim Biophys Acta. 2016; 1860: 1640–54.

22. Ogun OJ, Thaller G, Becker D. An Overview of the Importance and Value of Porcine Species in Sialic Acid Research. Biology (Basel). 2022; 11.

23. Mosquera JV, Auguste G, Wong D, Turner AW, Hodonsky CJ, Alvarez-Yela AC, et al. Integrative single-cell meta-analysis reveals disease-relevant vascular cell states and markers in human atherosclerosis. Cell Rep. 2023; 42: 113380.

24. Chen P-Y, Qin L, Li G, Malagon-Lopez J, Wang Z, Bergaya S, et al. Smooth Muscle Cell Reprogramming in Aortic Aneurysms. Cell Stem Cell. 2020; 26.

25. Wang M, Wang L, Pu L, Li K, Feng T, Zheng P, et al. LncRNAs related key pathways and genes in ischemic stroke by weighted gene co-expression network analysis (WGCNA). Genomics. 2020; 112: 2302–8.

26. Niu K, Bai P, Zhang J, Feng X, Qiu F. Cytidine Alleviates Dyslipidemia and Modulates the Gut Microbiota Composition in ob/ob Mice. Nutrients. 2023; 15.

27. Wang Y, Fan P, Zhang S, Wang L, Li X, Jia W, et al. Discrimination of Ribonucleoside Mono-, Di-, and Triphosphates Using an Engineered Nanopore. ACS Nano. 2022; 16: 21356–65.

28. Guertin DA, Wellen KE. Acetyl-CoA metabolism in cancer. Nat Rev Cancer. 2023; 23: 156–72.

29. Li X, Yu W, Qian X, Xia Y, Zheng Y, Lee J-H, et al. Nucleus-Translocated ACSS2 Promotes Gene Transcription for Lysosomal Biogenesis and Autophagy. Mol Cell. 2017; 66.

30. Xiang P, Chen Q, Chen L, Lei J, Yuan Z, Hu H, et al. Metabolite Neu5Ac triggers SLC3A2 degradation promoting vascular endothelial ferroptosis and aggravates atherosclerosis progression in ApoE-/-mice. Theranostics. 2023; 13: 4993–5016.

31. Zhang L, Wei T-T, Li Y, Li J, Fan Y, Huang F-Q, et al. Functional Metabolomics Characterizes a Key Role for N-Acetylneuraminic Acid in Coronary Artery Diseases. Circulation. 2018; 137: 1374–90.

32. Schiattarella GG, Wang Y, Tian R, Hill JA. Metabolism and Inflammation in Cardiovascular Health and Diseases: Mechanisms to Therapies. J Mol Cell Cardiol. 2021; 157: 113–4.

33. Chen L, Qiu H, Chen Q, Xiang P, Lei J, Zhang J, et al. N-acetylneuraminic acid modulates SQSTM1/p62 sialyation-mediated ubiquitination degradation contributing to vascular endothelium dysfunction in experimental atherosclerosis mice. IUBMB Life. 2023.

34. Li C, Zhao M, Xiao L, Wei H, Wen Z, Hu D, et al. Prognostic Value of Elevated Levels of Plasma N-Acetylneuraminic Acid in Patients With Heart Failure. Circ Heart Fail. 2021; 14: e008459.

35. Li M-N, Qian S-H, Yao Z-Y, Ming S-P, Shi X-J, Kang P-F, et al. Correlation of serum N-Acetylneuraminic acid with the risk and prognosis of acute coronary syndrome: a prospective cohort study. BMC Cardiovasc Disord. 2020; 20: 404.

36. Kawanishi K, Saha S, Diaz S, Vaill M, Sasmal A, Siddiqui SS, et al. Evolutionary conservation of human ketodeoxynonulosonic acid production is independent of sialoglycan biosynthesis. J Clin Invest. 2021; 131.

37. Lu Z, Ma H, Xu C, Shao Z, Cen C, Li Y. Serum Sialic Acid Level Is Significantly Associated with Nonalcoholic Fatty Liver Disease in a Nonobese Chinese Population: A Cross-Sectional Study. Biomed Res Int. 2016; 2016: 5921589.

38. Yildiz D, Ekin S, Sahinalp S. Evaluations of Antioxidant Enzyme Activities, Total Sialic Acid and Trace Element Levels in Coronary Artery Bypass Grafting Patients. Braz J Cardiovasc Surg. 2021; 36: 769–79.

39. Gokmen SS, Kilicli G, Ozcelik F, Ture M, Gulen S. Association between serum total and lipid-bound sialic acid concentration and the severity of coronary atherosclerosis. J Lab Clin Med. 2002; 140: 110–8.

40. Hu Z, Liu W, Hua X, Chen X, Chang Y, Hu Y, et al. Single-Cell Transcriptomic Atlas of Different Human Cardiac Arteries Identifies Cell Types Associated With Vascular Physiology. Arterioscler Thromb Vasc Biol. 2021; 41: 1408–27.

41. Xiong J, Li Z, Tang H, Duan Y, Ban X, Xu K, et al. Bulk and single-cell characterisation of the immune heterogeneity of atherosclerosis identifies novel targets for immunotherapy. BMC Biol. 2023; 21: 46.

42. Poznyak A, Grechko AV, Poggio P, Myasoedova VA, Alfieri V, Orekhov AN. The Diabetes Mellitus-Atherosclerosis Connection: The Role of Lipid and Glucose Metabolism and Chronic Inflammation. Int J Mol Sci. 2020; 21.

43. Neeland IJ, Poirier P, Després J-P. Cardiovascular and Metabolic Heterogeneity of Obesity: Clinical Challenges and Implications for Management. Circulation. 2018; 137: 1391–406.

44. Zernecke A, Erhard F, Weinberger T, Schulz C, Ley K, Saliba A-E, et al. Integrated single-cell analysis-based classification of vascular mononuclear phagocytes in mouse and human atherosclerosis. Cardiovasc Res. 2023; 119: 1676–89.

45. Wang Y, Yang H, Geerts C, Furtos A, Waters P, Cyr D, et al. The multiple facets of acetyl-CoA metabolism: Energetics, biosynthesis, regulation, acylation and inborn errors. Mol Genet Metab. 2023; 138: 106966.

46. Ritterhoff J, Young S, Villet O, Shao D, Neto FC, Bettcher LF, et al. Metabolic Remodeling Promotes Cardiac Hypertrophy by Directing Glucose to Aspartate Biosynthesis. Circ Res. 2020; 126: 182–96.

47. Poznyak AV, Kashirskikh DA, Postnov AY, Popov MA, Sukhorukov VN, Orekhov AN. Sialic acid as the potential link between lipid metabolism and inflammation in the pathogenesis of atherosclerosis. Braz J Med Biol Res. 2023; 56: e12972.

48. Hu X, Li Y, Chen Q, Wang T, Ma L, Zhang W, et al. Sialic acids promote macrophage M1 polarization and atherosclerosis by upregulating ROS and autophagy blockage. Int Immunopharmacol. 2023; 120: 110410.

49. Chen W, Schilperoort M, Cao Y, Shi J, Tabas I, Tao W. Macrophage-targeted nanomedicine for the diagnosis and treatment of atherosclerosis. Nat Rev Cardiol. 2022; 19: 228–49.

50. Sorto P, Mäyränpää MI, Saksi J, Nuotio K, Ijäs P, Tuimala J, et al. Glutamine synthetase in human carotid plaque macrophages associates with features of plaque vulnerability: An immunohistological study. Atherosclerosis. 2022; 352: 18–26.

51. Chen Y, Wu G, Li M, Hesse M, Ma Y, Chen W, et al. LDHA-mediated metabolic reprogramming promoted cardiomyocyte proliferation by alleviating ROS and inducing M2 macrophage polarization. Redox Biol. 2022; 56: 102446.

52. Lin Y, Zhang W, Liu L, Li W, Li Y, Li B. ENO1 Promotes OSCC Migration and Invasion by Orchestrating IL-6 Secretion from Macrophages via a Positive Feedback Loop. Int J Mol Sci. 2023; 24.

53. Zhang Y, Yu G, Chu H, Wang X, Xiong L, Cai G, et al. Macrophage-Associated PGK1 Phosphorylation Promotes Aerobic Glycolysis and Tumorigenesis. Mol Cell. 2018; 71.

54. Zhou X, Zhang C, Yang S, Yang L, Luo W, Zhang W, et al. Macrophage-derived MMP12 promotes fibrosis through sustained damage to ECs. J Hazard Mater. 2024; 461: 132733.

55. Ke K, Wu Z, Lin J, Lin L, Huang N, Yang W. Increased Expression of CD74 in Atherosclerosis Associated with Inflammatory Responses of ECs and Macrophages. Biochem Genet. 2023.

56. Chen Q, Guo C, Zhou X, Su Y, Guo H, Cao M, et al. N-acetylneuraminic acid and chondroitin sulfate modified nanomicelles with ROS-sensitive H2S donor via targeting E-selectin receptor and CD44 receptor for the efficient therapy of atherosclerosis. Int J Biol Macromol. 2022; 211: 259–70.

