## supplementary figures and tables for "Single-Cell Transcriptomic Profiling Unveils Critical Metabolic Alterations and Signatures in Progression of Atherosclerosis": Supplement Figures.docx

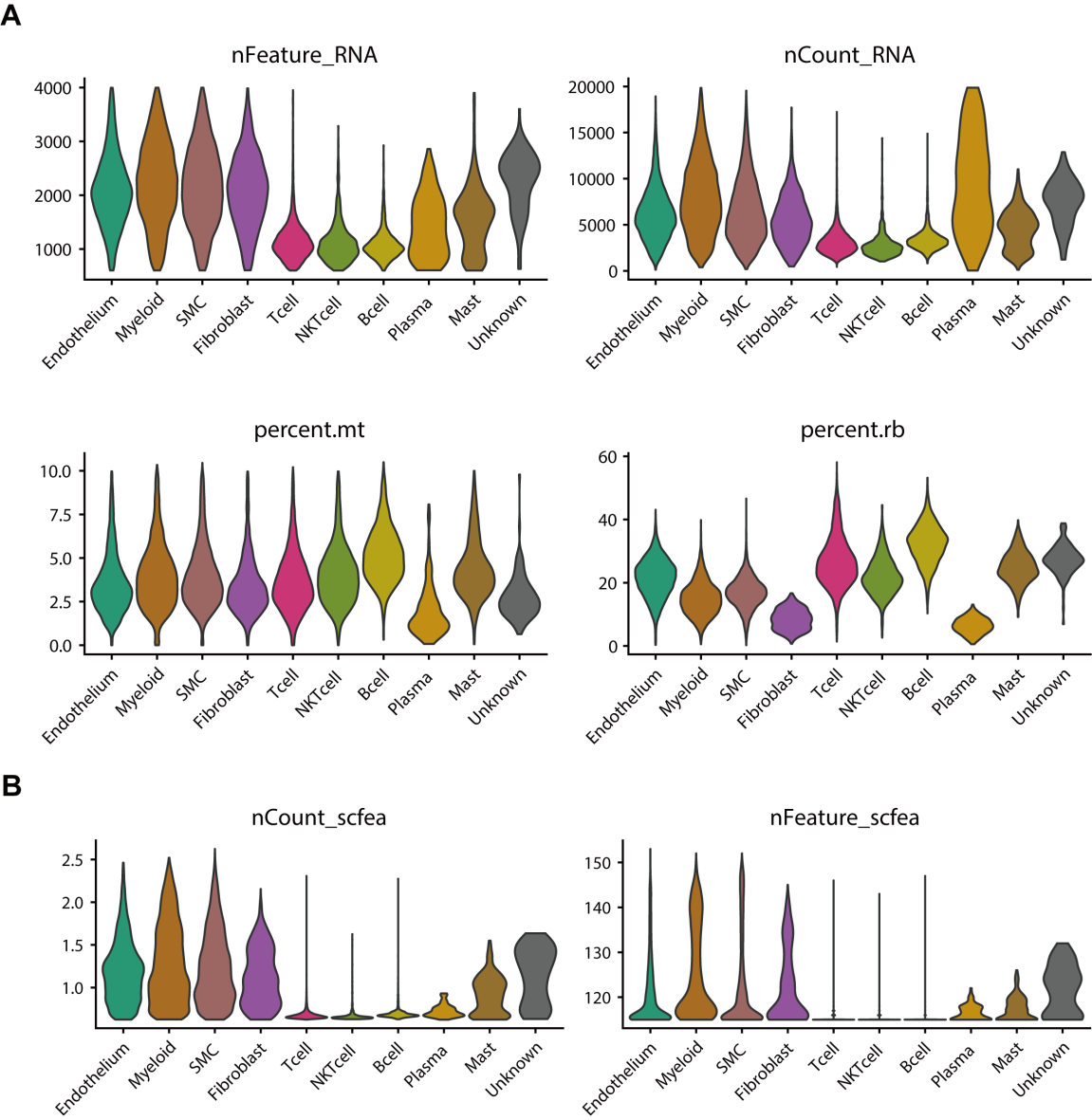

Supplementary Fig. 1 Quality control and filtering of cells in Single-cell levels. (A) Violin plots showing the cells quality control in steps like nFeature-RNA, nCount-RNA, percent.me and percent rb, and then used for the cellular heterogeneity analysis. (B) Violin plots showing the quality control and filtering of cells for scFFA analysis.

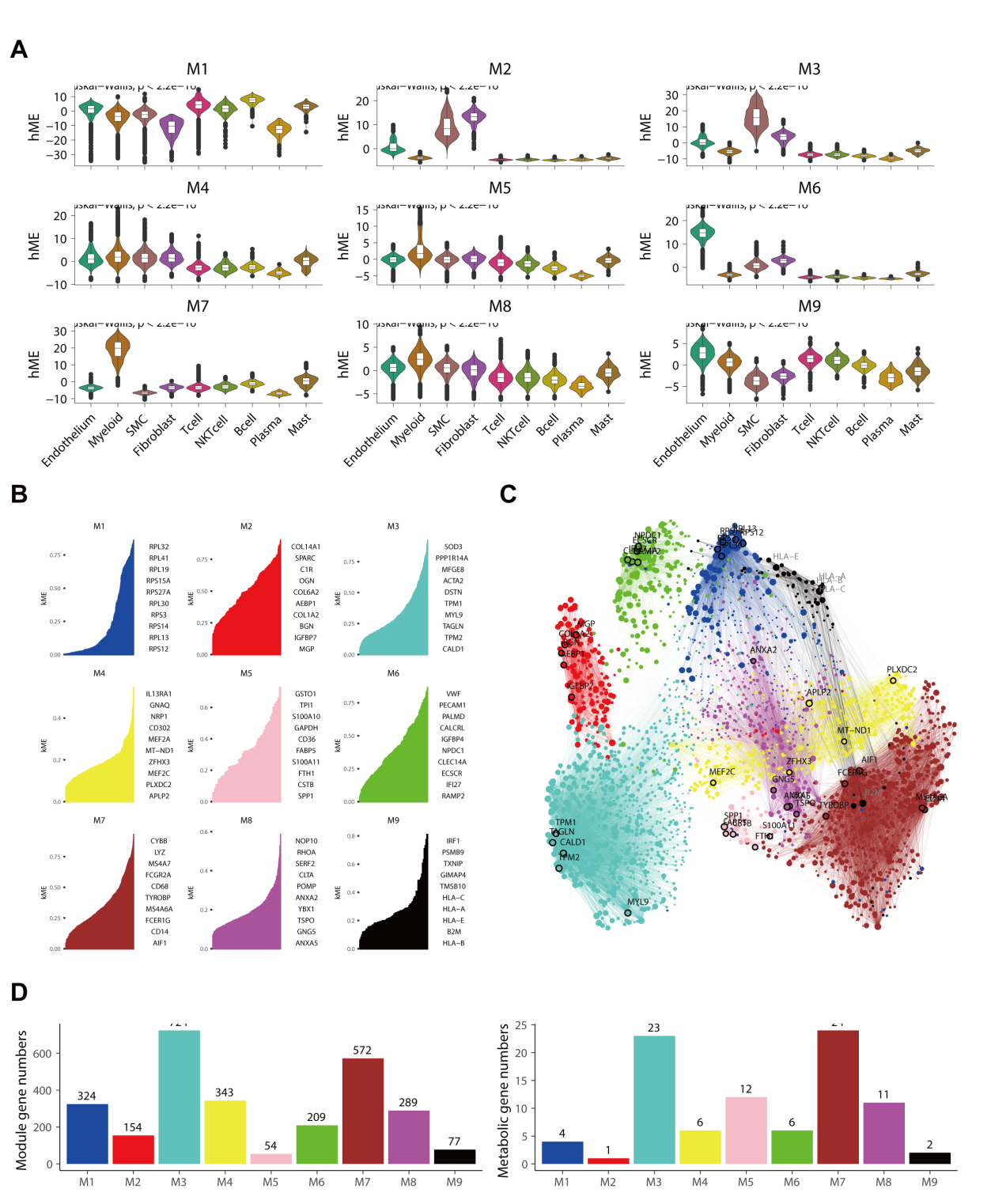

Supplementary Fig. 2 scWGCNA analysis revealed the different modules and metabolic shift in different cell types. (A) Profiles of the predicted modules in different cell types. (B) Identification of hub genes in different modules (M1-M9). (C) Cell-cell interaction analysis between modules and the color showing the corresponding modules, highlighted the genes which play an important role in regulating interaction. (D) Gene numbers in different modules and the metabolic genes which differs in AC vs PA group in different modules.

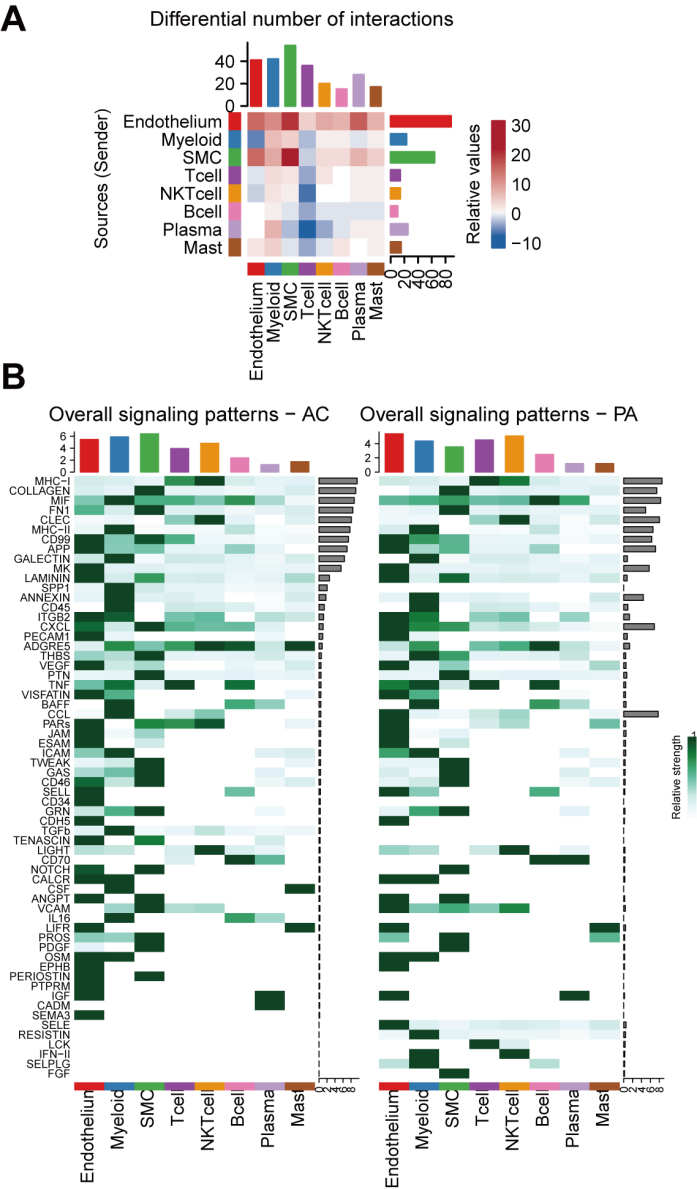

Supplementary Fig. 3 Cell-cell interaction analysis revealed the potential communication in AS progression. (A) Relative value of differential interaction in various cell types. The red color indicated that the interaction was enhanced while the blue color indicated the interaction displayed decreasing. (B) Comparison of overall signaling patterns of cells between the PA and AC groups. The color is proportional to the contribution score computed from pattern recognition analysis. A higher score implies that the signaling pathway is more enriched in the corresponding cell group.
